## Supplemental Information for "Cerebrovascular reactivity assessment with O_2_-CO_2_ exchange ratio under brief breath hold challenge"

### Supporting information

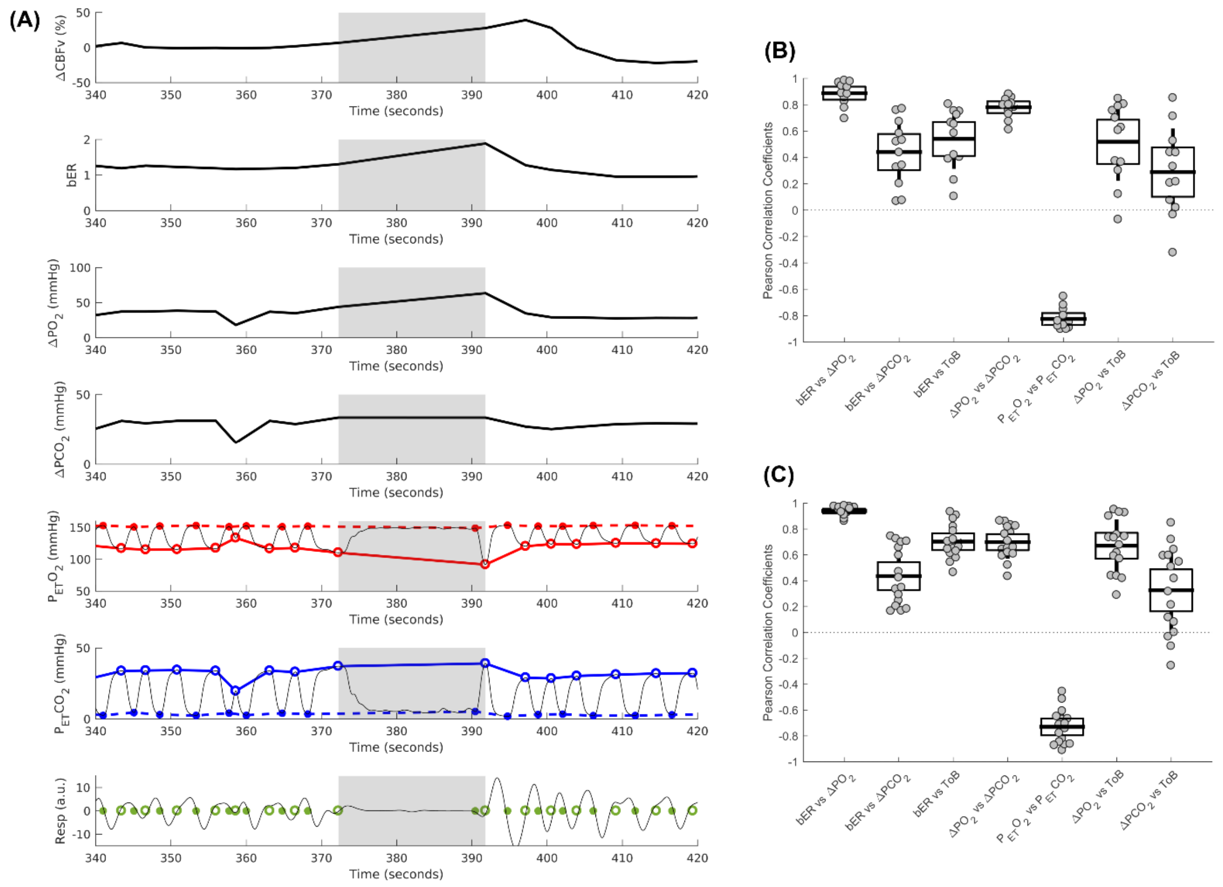


**S1 Fig. Definition of end inspiration and end expiration on the time series of RGE metrics and the correlations among breath-by breath RGE matrices.** (A) A segment of 80-second time series of ΔCBFv in left MCA and physiological changes including breath-by-breath bER, $\Delta PO_2$, $\Delta PCO_2$, $P_{ET}O_2$ and $P_{ET}CO_2$ measured by gas analyzers and respiration time series (Resp) measured by respiratory bellow in a representative subject under breath hold challenge in TCD session. Open circles represent end expiration while closed circles represent end inspiration in resting phase or onset of expiration at the end of breath hold epoch. Positive phases with deflection above zero on the respiration time series represent inspiration and negative phases with deflection below zero represent expiration. The inspiratory and expiratory phases of each respiratory cycle on the time series of $P_{ET}O_2$ and $P_{ET}CO_2$ are verified by those on respiration time series. The timing for open (end expiration) and closed (end inspiration) circles in green is the same as those in red and blue. (B) Correlations among breath-by breath respiratory matrices (bER, $\Delta PO_2$, $\Delta PCO_2$, ToB, $P_{ET}O_2$ and $P_{ET}CO_2$) in all subjects who participated in TCD sessions (n=12), and (C) those who participated in MRI sessions (n=16). Each gray circle represents the Pearson's correlation coefficient from the correlation analysis of the time series of parameter pair shown on x-axis for each subject. The thick middle horizontal line, the box and the vertical rod represent the mean, standard deviation and 95% confidence interval of the group data respectively. The time series of bER had stronger correlation with that of $\Delta PO_2$ than $\Delta PCO_2$, although both $\Delta PO_2$ and $\Delta PCO_2$ contributed to changes of bER. The correlation coefficients from $\Delta PO_2$ vs $\Delta PCO_2$ varied from 0.6 to 0.9 in TCD sessions and from 0.4 to 0.9 in MRI sessions, suggesting that $\Delta PO_2$ and $\Delta PCO_2$ are not necessarily redundant. The difference in the ranges of correlation strength found between TCD and MRI sessions may be due to the difference in posture of the subjects, where the subjects were in erect seated position in TCD sessions and they were in supine position in MRI sessions.

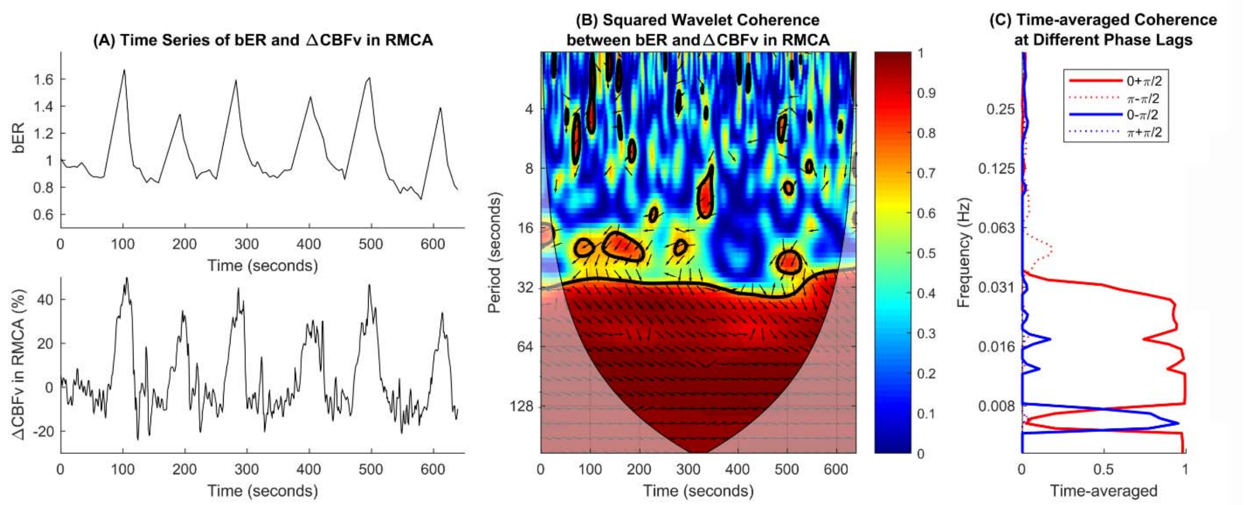


**S2 Fig. Wavelet transform coherence analysis between bER and ΔCBFv in a representative subject.** (A) Time series of bER and ΔCBFv measured in right MCA in a representative subject under breath hold challenge. (B) The squared wavelet coherence between these two time series. Squared wavelet coherence is plotted with x-axis as time and y-axis as scale which has been converted to its equivalent Fourier period. The magnitude of wavelet transform coherence ranges between 0 and 1, where warmer color represents stronger coherence and cooler color represents weaker coherence. Areas inside the 'cone of influence', which are locations in the time-frequency plane where edge effects give rise to lower confidence in the computed values, are shown in faded color outside of the conical contour. The statistical significance level of the wavelet coherence is estimated using Monte Carlo methods and the 5% significance level against red noise is shown as thick contour. The phase angle between the two time series at particular samples of the time-frequency plane is indicated by an arrow (rightward pointing arrows indicate that the time series are in phase or positively correlation, leftward pointing arrows indicate anticorrelation and the downward pointing arrows indicate phase angles of π/2). There are four different ranges of phase lags: 0+π/2, 0-π/2, π-π/2, and π+π/2. (C) Time-averaged coherences at four different phase lags of 0+π/2, 0-π/2, π-π/2, and π+π/2. At each phase lag range, time-averaged coherence was defined as the total significant coherence at each scale where the wavelet coherence magnitude exceeded 95% significance level, normalized by the maximum possible coherence outside the cone of influence, i.e. inside the conical contour, at that particular scale and phase lag range.

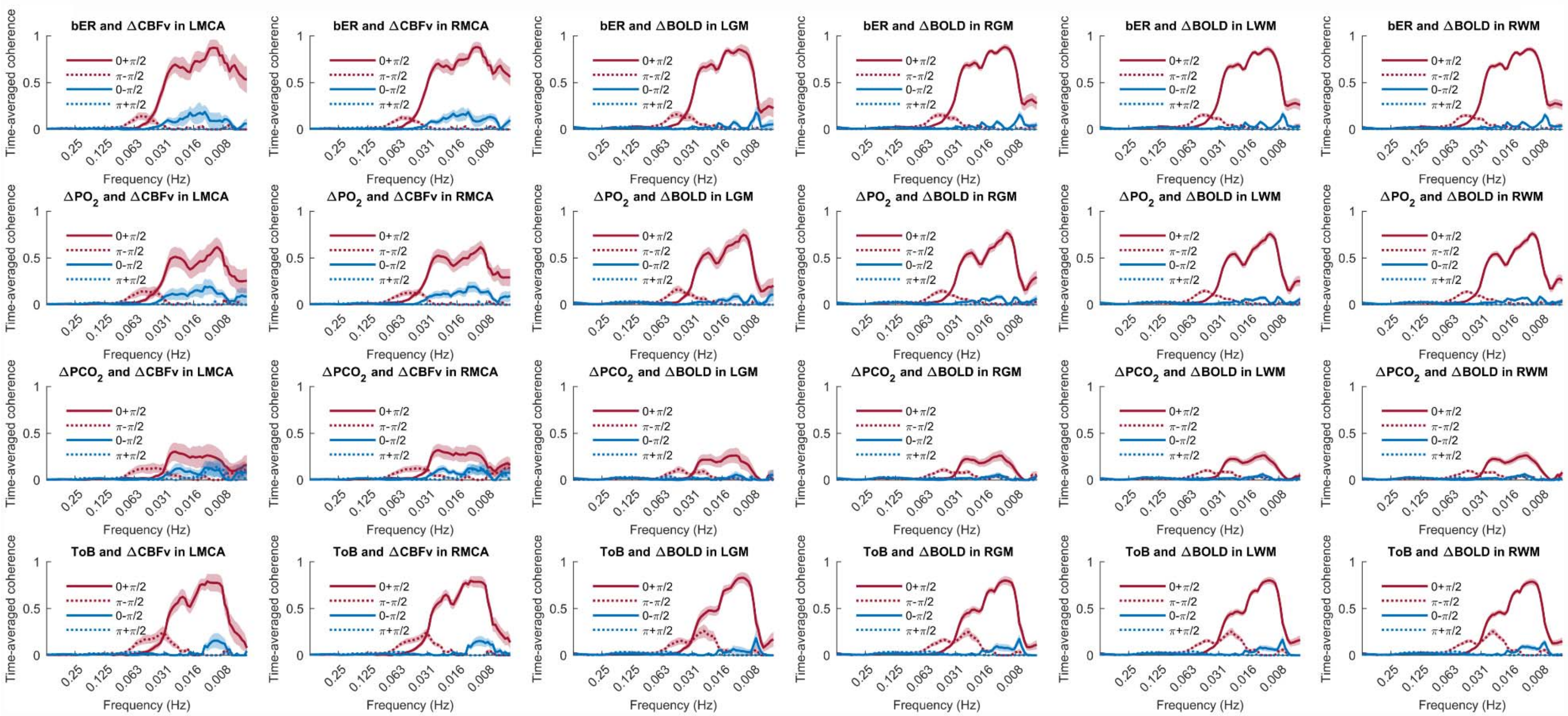


**S3 Fig. Coherence between time series of RGE metrics and cerebral hemodynamic response at four different phase lags (0+π/2, 0-π/2, π-π/2, and π+π/2).** The mean time-averaged coherence between time series of respiratory metrics and cerebral hemodynamic responses (ΔCBFv in LMCA and RMCA, and ΔBOLD in LGM, RGM, LWM and RWM) at four different phase lags (0+π/2, 0-π/2, π-π/2, and π+π/2) for the subjects included in the TCD sessions (n=12) and in the MRI sessions (n=16). Color shaded areas represent SEM. Comparing with $\Delta PO_2$, $\Delta PCO_2$ and ToB, the total time-averaged coherence between bER and cerebral hemodynamic responses was found to be significantly stronger between 0.008Hz (1/128 seconds) and 0.03Hz (1/32 seconds). The strong mean time-averaged coherence between respiratory metrics and cerebral hemodynamic responses were found at phase lag of $0+\pi/2$.

**S1 Table. Correlation among RGE metrics.**

| Subjects | bER vs $\Delta PO_2$ | bER vs $\Delta PCO_2$ | bER vs ToB | $\Delta PO_2$ vs $\Delta PCO_2$ | $P_{ET}O_2$ vs $P_{ET}CO_2$ | $\Delta PO_2$ vs ToB | $\Delta PCO_2$ vs ToB |
|---|---|---|---|---|---|---|---|
| | | | | ***TCD sessions*** | | | |
| s4 | 0.777* | 0.326* | 0.389* | 0.840* | -0.878* | 0.374* | 0.207□ |
| s5 | 0.888* | 0.341* | 0.420* | 0.730* | -0.799* | 0.365* | 0.076 |
| s6 | 0.916* | 0.520* | 0.665* | 0.799* | -0.840* | 0.700* | 0.439* |
| s7 | 0.969* | 0.672* | 0.653* | 0.826* | -0.870* | 0.628* | 0.332* |
| s8 | 0.944* | 0.583* | 0.725* | 0.811* | -0.849* | 0.756* | 0.526* |
| s9 | 0.883* | 0.438* | 0.105 | 0.802* | -0.849* | 0.122 | 0.019 |
| s10 | 0.934* | 0.532* | 0.805* | 0.783* | -0.902* | 0.788* | 0.437* |
| s11 | 0.984* | 0.759* | 0.753* | 0.857* | -0.892* | 0.805* | 0.713* |
| s12 | 0.977* | 0.771* | 0.751* | 0.881* | -0.902* | 0.847* | 0.852* |
| s14 | 0.696* | 0.068 | 0.229† | 0.757* | -0.719* | -0.070 | -0.322* |
| s15 | 0.855* | 0.203‡ | 0.403* | 0.673* | -0.654* | 0.302* | -0.034 |
| s17 | 0.827* | 0.075 | 0.585* | 0.612* | -0.752* | 0.606* | 0.216‡ |
| | | | | ***BOLD sessions*** | | | |
| s1 | 0.887* | 0.166 | 0.613* | 0.595* | -0.649* | 0.439* | -0.106 |
| s2 | 0.976* | 0.745* | 0.819* | 0.864* | -0.873* | 0.901* | 0.849* |
| s3 | 0.953* | 0.471* | 0.934* | 0.709* | -0.778* | 0.951* | 0.594* |
| s4 | 0.862* | 0.336* | 0.645* | 0.762* | -0.820* | 0.438* | -0.032 |
| s5 | 0.963* | 0.658* | 0.629* | 0.818* | -0.846* | 0.617* | 0.317□ |
| s6 | 0.969* | 0.726* | 0.704* | 0.860* | -0.911* | 0.757* | 0.597* |
| s7 | 0.972* | 0.425* | 0.881* | 0.619* | -0.607* | 0.931* | 0.642* |
| s8 | 0.914* | 0.247‡ | 0.785* | 0.608* | -0.672* | 0.736* | 0.214‡ |
| s9 | 0.975* | 0.700* | 0.722* | 0.833* | -0.867* | 0.734* | 0.509* |
| s10 | 0.980* | 0.706* | 0.907* | 0.825* | -0.863* | 0.928* | 0.720* |
| s11 | 0.880* | 0.204‡ | 0.544* | 0.642* | -0.667* | 0.290* | -0.256□ |
| s12 | 0.958* | 0.168 | 0.648* | 0.435* | -0.514* | 0.592* | 0.002 |
| s13 | 0.909* | 0.293□ | 0.465* | 0.654* | -0.712* | 0.419* | 0.116 |
| s14 | 0.932* | 0.183 | 0.579* | 0.521* | -0.456* | 0.678* | 0.418* |
| s15 | 0.968* | 0.597* | 0.697* | 0.773* | -0.742* | 0.743* | 0.547* |
| s16 | 0.928* | 0.349* | 0.663* | 0.660* | -0.705* | 0.571* | 0.081 |

**p≤0.001, ⁑p≤0.005, †p≤0.01, ‡p≤0.05*

Strength of correlation indicated by Pearson's correlation coefficients among respiratory metrics including bER, $\Delta PO_2$, $\Delta PCO_2$, ToB, $P_{ET}O_2$ and $P_{ET}CO_2$ in all subjects who participated in TCD sessions (n=12), and those who participated in MRI sessions (n=16). The time series of bER had stronger correlation with that of $\Delta PO_2$ than $\Delta PCO_2$, although both $\Delta PO_2$ and $\Delta PCO_2$ contributed to changes of bER. The correlation coefficients from $\Delta PO_2$ vs $\Delta PCO_2$ varied from 0.6 to 0.9 in TCD sessions and from 0.4 to 0.9 in MRI sessions, suggesting that $\Delta PO_2$ than $\Delta PCO_2$ are not necessarily redundant.

**S2 Table A. Correlation between RGE metrics and ΔCBFv in TCD sessions.**

| | ΔCBFv in LMCA | | | | ΔCBFv in RMCA | | | |
|---|---|---|---|---|---|---|---|---|
| **Subjects** | **bER** | **$\Delta PO_2$** | **$\Delta PCO_2$** | **ToB** | **bER** | **$\Delta PO_2$** | **$\Delta PCO_2$** | **ToB** |
| s4 | 0.819 (<0.001) | 0.687 (<0.001) | 0.349 (<0.001) | 0.404 (<0.001) | 0.813 (<0.001) | 0.674 (<0.001) | 0.338 (<0.001) | 0.433 (<0.001) |
| s5 | 0.712 (<0.001) | 0.575 (<0.001) | 0.158 (0.122) | 0.318 (0.002) | 0.813 (<0.001) | 0.668 (<0.001) | 0.172 (0.092) | 0.491 (<0.001) |
| s6 | 0.838 (<0.001) | 0.685 (<0.001) | 0.264 (0.016) | 0.462 (<0.001) | 0.842 (<0.001) | 0.705 (<0.001) | 0.302 (0.006) | 0.460 (<0.001) |
| s7 | 0.754 (<0.001) | 0.645 (<0.001) | 0.302 (0.002) | 0.511 (<0.001) | 0.717 (<0.001) | 0.626 (<0.001) | 0.331 (0.001) | 0.486 (<0.001) |
| s8 | 0.733 (<0.001) | 0.556 (<0.001) | 0.072 (0.399) | 0.410 (<0.001) | 0.719 (<0.001) | 0.548 (<0.001) | 0.072 (0.400) | 0.421 (<0.001) |
| s9 | 0.433 (<0.001) | 0.277 (0.023) | -0.043 (0.730) | 0.349 (0.004) | 0.396 (0.001) | 0.247 (0.044) | -0.053 (0.670) | 0.345 (0.004) |
| s10 | 0.779 (<0.001) | 0.703 (<0.001) | 0.367 (<0.001) | 0.551 (<0.001) | 0.773 (<0.001) | 0.695 (<0.001) | 0.353 (<0.001) | 0.591 (<0.001) |
| s11 | --- | --- | --- | --- | 0.887 (<0.001) | 0.825 (<0.001) | 0.521 (<0.001) | 0.545 (<0.001) |
| s12 | 0.857 (<0.001) | 0.767 (<0.001) | 0.482 (<0.001) | 0.389 (<0.001) | 0.865 (<0.001) | 0.767 (<0.001) | 0.453 (<0.001) | 0.388 (<0.001) |
| s14 | 0.813 (<0.001) | 0.440 (<0.001) | -0.159 (0.074) | 0.269 (0.002) | 0.810 (<0.001) | 0.433 (<0.001) | -0.165 (0.064) | 0.253 (0.004) |
| s15 | 0.837 (<0.001) | 0.669 (<0.001) | 0.058 (0.496) | 0.468 (<0.001) | 0.828 (<0.001) | 0.661 (<0.001) | 0.058 (0.491) | 0.448 (<0.001) |
| s17 | 0.652 (<0.001) | 0.400 (<0.001) | -0.201 (0.019) | 0.372 (<0.001) | 0.684 (<0.001) | 0.429 (<0.001) | -0.194 (0.024) | 0.384 (<0.001) |
| Mean Fisher Z | 1.008 (---) | 0.691 (<0.001) | 0.158 (<0.001) | 0.439 (<0.001) | 1.052 (---) | 0.739 (<0.001) | 0.194 (<0.001) | 0.474 (<0.001) |

Strength of correlation indicated by Pearson's correlation coefficients between ΔCBFv and RGE metrics including bER, $\Delta PO_2$, $\Delta PCO_2$ and ToB (n=12). Numbers in brackets next to Pearson's correlation coefficients indicate p values from individual correlation analyses. The bottom row shows the mean values of Fisher Z scores transformed from Pearson's correlation coefficients in groups. Numbers in brackets next to mean Fisher Z scores indicate p values in the paired comparisons. The correlation between ΔCBFv and bER was significantly larger than those of the correlation between ΔCBFv and the other respiratory metrics in the paired comparisons ($p<0.001$). bER is the only parameter that consistently showed significantly high correlation with the ΔCBFv measured in LMCA and RMCA.

**S2 Table B. Correlation between RGE metrics and $\Delta$BOLD in MRI sessions.**

| | $\Delta$BOLD in LGM | | | | $\Delta$BOLD in RGM | | | |
|---|---|---|---|---|---|---|---|---|
| Subjects | bER | $\Delta PO_2$ | $\Delta PCO_2$ | ToB | bER | $\Delta PO_2$ | $\Delta PCO_2$ | ToB |
| s1 | 0.674 (<0.001) | 0.558 (<0.001) | 0.044 (0.653) | 0.274 (0.005) | 0.696 (<0.001) | 0.577 (<0.001) | 0.044 (0.656) | 0.304 (0.002) |
| s2 | 0.810 (<0.001) | 0.744 (<0.001) | 0.433 (<0.001) | 0.616 (<0.001) | 0.829 (<0.001) | 0.758 (<0.001) | 0.436 (<0.001) | 0.631 (<0.001) |
| s3 | 0.511 (<0.001) | 0.472 (<0.001) | 0.160 (0.252) | 0.494 (<0.001) | 0.498 (<0.001) | 0.445 (0.001) | 0.109 (0.436) | 0.463 (<0.001) |
| s4 | 0.795 (<0.001) | 0.658 (<0.001) | 0.190 (0.030) | 0.447 (<0.001) | 0.762 (<0.001) | 0.618 (<0.001) | 0.157 (0.075) | 0.435 (<0.001) |
| s5 | 0.352 (0.001) | 0.344 (0.002) | 0.185 (0.101) | 0.410 (<0.001) | 0.579 (<0.001) | 0.555 (<0.001) | 0.352 (0.001) | 0.510 (<0.001) |
| s6 | 0.535 (<0.001) | 0.419 (<0.001) | 0.100 (0.357) | 0.250 (0.020) | 0.576 (<0.001) | 0.452 (<0.001) | 0.118 (0.275) | 0.265 (0.013) |
| s7 | 0.723 (<0.001) | 0.662 (<0.001) | 0.133 (0.204) | 0.490 (<0.001) | 0.786 (<0.001) | 0.742 (<0.001) | 0.234 (0.024) | 0.582 (<0.001) |
| s8 | 0.708 (<0.001) | 0.556 (<0.001) | -0.070 (0.483) | 0.374 (<0.001) | 0.733 (<0.001) | 0.595 (<0.001) | -0.022 (0.826) | 0.398 (<0.001) |
| s9 | 0.673 (<0.001) | 0.582 (<0.001) | 0.233 (0.075) | 0.621 (<0.001) | 0.675 (<0.001) | 0.587 (<0.001) | 0.244 (0.063) | 0.640 (<0.001) |
| s10 | 0.605 (<0.001) | 0.519 (<0.001) | 0.220 (0.026) | 0.469 (<0.001) | 0.735 (<0.001) | 0.659 (<0.001) | 0.366 (<0.001) | 0.576 (<0.001) |
| s11 | 0.175 (0.056) | -0.067 (0.466) | -0.423 (<0.001) | 0.367 (<0.001) | 0.311 (0.001) | 0.069 (0.452) | -0.347 (<0.001) | 0.335 (<0.001) |
| s12 | 0.698 (<0.001) | 0.629 (<0.001) | -0.057 (0.543) | 0.298 (0.001) | 0.672 (<0.001) | 0.601 (<0.001) | -0.063 (0.497) | 0.259 (0.005) |
| s13 | 0.478 (<0.001) | 0.396 (<0.001) | 0.066 (0.528) | 0.298 (0.004) | 0.474 (<0.001) | 0.411 (<0.001) | 0.109 (0.300) | 0.297 (0.004) |
| s14 | 0.614 (<0.001) | 0.551 (<0.001) | 0.047 (0.646) | 0.371 (<0.001) | 0.707 (<0.001) | 0.652 (<0.001) | 0.113 (0.265) | 0.369 (<0.001) |
| s15 | 0.717 (<0.001) | 0.631 (<0.001) | 0.283 (0.005) | 0.375 (<0.001) | 0.414 (<0.001) | 0.355 (<0.001) | 0.146 (0.155) | 0.040 (0.696) |
| s16 | 0.525 (<0.001) | 0.423 (<0.001) | -0.032 (0.775) | 0.304 (0.006) | 0.564 (<0.001) | 0.461 (<0.001) | -0.021 (0.855) | 0.337 (0.002) |
| Mean Fisher Z | 0.727 (---) | 0.579 (<0.001) | 0.096 (<0.001) | 0.436 (<0.001) | 0.767 (---) | 0.620 (<0.001) | 0.127 (<0.001) | 0.441 (<0.001) |

| | $\Delta$BOLD in LWM | | | | $\Delta$BOLD in RWM | | | |
|---|---|---|---|---|---|---|---|---|
| Subjects | bER | $\Delta PO_2$ | $\Delta PCO_2$ | ToB | bER | $\Delta PO_2$ | $\Delta PCO_2$ | ToB |
| s1 | 0.648 (<0.001) | 0.540 (<0.001) | 0.057 (0.560) | 0.268 (0.005) | 0.655 (<0.001) | 0.547 (<0.001) | 0.052 (0.597) | 0.287 (0.003) |
| s2 | 0.792 (<0.001) | 0.740 (<0.001) | 0.472 (<0.001) | 0.626 (<0.001) | 0.806 (<0.001) | 0.741 (<0.001) | 0.451 (<0.001) | 0.621 (<0.001) |
| s3 | 0.543 (<0.001) | 0.480 (<0.001) | 0.096 (0.496) | 0.538 (<0.001) | 0.464 (<0.001) | 0.392 (0.004) | 0.030 (0.833) | 0.419 (0.002) |
| s4 | 0.789 (<0.001) | 0.654 (<0.001) | 0.193 (0.028) | 0.408 (<0.001) | 0.770 (<0.001) | 0.627 (<0.001) | 0.164 (0.062) | 0.428 (<0.001) |
| s5 | 0.403 (<0.001) | 0.394 (<0.001) | 0.237 (0.034) | 0.434 (<0.001) | 0.588 (<0.001) | 0.556 (<0.001) | 0.328 (0.003) | 0.584 (<0.001) |
| s6 | 0.571 (<0.001) | 0.454 (<0.001) | 0.132 (0.223) | 0.262 (0.014) | 0.589 (<0.001) | 0.466 (<0.001) | 0.134 (0.216) | 0.265 (0.013) |
| s7 | 0.548 (<0.001) | 0.497 (<0.001) | 0.081 (0.442) | 0.353 (0.001) | 0.746 (<0.001) | 0.700 (<0.001) | 0.218 (0.036) | 0.515 (<0.001) |
| s8 | 0.702 (<0.001) | 0.558 (<0.001) | -0.053 (0.594) | 0.378 (<0.001) | 0.749 (<0.001) | 0.605 (<0.001) | -0.027 (0.784) | 0.423 (<0.001) |
| s9 | 0.642 (<0.001) | 0.551 (<0.001) | 0.214 (0.103) | 0.591 (<0.001) | 0.679 (<0.001) | 0.585 (<0.001) | 0.232 (0.077) | 0.580 (<0.001) |
| s10 | 0.620 (<0.001) | 0.538 (<0.001) | 0.245 (0.013) | 0.487 (<0.001) | 0.722 (<0.001) | 0.651 (<0.001) | 0.382 (<0.001) | 0.569 (<0.001) |
| s11 | -0.011 (0.902) | -0.184 (0.044) | -0.374 (<0.001) | 0.295 (0.001) | 0.213 (0.019) | 0.043 (0.644) | -0.243 (0.008) | 0.226 (0.013) |

| | | | | | | | | |
|---|---|---|---|---|---|---|---|---|
| s12 | 0.634 (<0.001) | 0.571 (<0.001) | -0.043 (0.645) | 0.234 (0.011) | 0.662 (<0.001) | 0.583 (<0.001) | -0.096 (0.303) | 0.253 (0.006) |
| s13 | 0.262 (0.011) | 0.217 (0.037) | 0.039 (0.708) | 0.064 (0.542) | 0.257 (0.013) | 0.241 (0.020) | 0.103 (0.327) | 0.028 (0.793) |
| s14 | 0.581 (<0.001) | 0.569 (<0.001) | 0.174 (0.086) | 0.402 (<0.001) | 0.530 (<0.001) | 0.493 (<0.001) | 0.111 (0.275) | 0.222 (0.027) |
| s15 | 0.742 (<0.001) | 0.656 (<0.001) | 0.308 (0.002) | 0.379 (<0.001) | 0.609 (<0.001) | 0.530 (<0.001) | 0.232 (0.022) | 0.170 (0.097) |
| s16 | 0.500 (<0.001) | 0.405 (<0.001) | -0.025 (0.827) | 0.262 (0.018) | 0.540 (<0.001) | 0.436 (<0.001) | -0.032 (0.775) | 0.310 (0.005) |
| Mean Fisher Z | 0.672 (---) | 0.544 (<0.001) | 0.113 (<0.001) | 0.403 (0.001) | 0.726 (---) | 0.589 (<0.001) | 0.132 (<0.001) | 0.402 (<0.001) |

Strength of correlation indicated by Pearson's correlation coefficients between ΔBOLD and RGE metrics including bER, $\Delta PO_2$, $\Delta PCO_2$ and ToB (n=16). Numbers in brackets next to Pearson's correlation coefficients indicate p values from individual correlation analyses. The bottom row shows the mean values of Fisher Z scores transformed from Pearson's correlation coefficients in groups. Numbers in brackets next to mean Fisher Z scores indicate p values in paired comparisons. The correlation between ΔBOLD and bER was significantly larger than those of the correlation between ΔBOLD and the other respiratory metrics in the paired comparisons ($p<0.001$). bER is the only parameter that consistently showed significantly high correlation with the ΔBOLD measured in LGM, RGM, LWM and RWM.
